# Cell Painting, an image-based assay for morphological profiling

**DOI:** 10.1101/049817

**Authors:** Mark-Anthony Bray, Shantanu Singh, Han Han, Chadwick T. Davis, Blake Borgeson, Cathy Hartland, Maria Kost-Alimova, Sigrun M. Gustafsdottir, Christopher C. Gibson, Anne E. Carpenter

## Abstract

In morphological profiling, quantitative data are extracted from microscopy images of cells to identify biologically relevant similarities and differences among samples based on these profiles. This protocol describes the design and execution of experiments using Cell Painting, a morphological profiling assay multiplexing six fluorescent dyes imaged in five channels, to reveal eight broadly relevant cellular components or organelles. Automated image analysis software identifies individual cells and measures ~1,500 morphological features (various measures of size, shape, texture, intensity, etc.) to produce a rich profile suitable for detecting subtle phenotypes. Profiles of cell populations treated with different experimental perturbations can be compared to suit many goals, such as identifying the phenotypic impact of chemical or genetic perturbations, grouping compounds and/or genes into functional pathways, and identifying signatures of disease. Cell culture and image acquisition takes 2 weeks; feature extraction and data analysis take an additional 1-2 weeks.

## INTRODUCTION

Phenotypic screening has been tremendously powerful for identifying novel small molecules as probes and potential therapeutics and for identifying genetic regulators of many biological processes^1–4^. High-throughput microscopy has been a particularly fruitful type of phenotypic screening; it is often called high-content analysis because of the potentially high information content that can be observed in images^5^. However, most large-scale imaging experiments extract only one or two features of cells^6^ and/or aim to identify just a few “hits” in a screen, meaning that vast quantities of quantitative data about cellular state remain unharnessed.

In this article, we detail a protocol for the Cell Painting assay, a generalizable and broadly-applicable method for accessing the valuable biological information about cellular state that is contained in morphology. Cellular morphology is a potentially rich data source as a means of interrogating biological perturbations, especially in large scale^5,7–10^. The techniques and technology necessary to generate these data have advanced rapidly, and are now becoming accessible to non-specialized laboratories^11^. In this protocol, we discuss morphological profiling (also known as image-based profiling), contrast it with conventional image-based screening, illustrate applications of morphological profiling, and provide guidance, tips, and tricks related to the successful execution of one particular morphological profiling assay, the Cell Painting assay.

Broadly speaking, the term *profiling* describes the process of quantifying a very large set of features, typically hundreds to thousands, from each experimental sample in a relatively unbiased way. Significant changes in a subset of profiled features can thus serve as a “fingerprint” characterizing the sample condition. Some of the earliest instances of profiling involved the NCI-60 tumor cell line panel, where patterns of anticancer drug sensitivity were discovered to reflect mechanisms of action^12^, and gene expression, in which signatures related to small molecules, genes, and diseases were identified^13^.

It is important to note that profiling differs from conventional screening assays in that the latter are focused on quantifying a relatively small number of features selected specifically because of a known association with the biology of interest. Profiling, on the other hand, casts a much wider net and avoids the intensive customization usually necessary for problem-specific assay development in favor of a more generalizable method. Therefore, taking an unbiased approach via morphological profiling offers the opportunity for discovery unconstrained by what we know (or think we know). It also holds the potential to be more efficient, as a single experiment can be mined for many different biological processes or diseases of interest.

In morphological profiling, measured features include staining intensities, textural patterns, size, and shape of the labeled cellular structures, as well as correlation between stains across channels and neighborhood relationships between cells and among intracellular structures. The technique enables single-cell resolution, enabling detection of perturbations even in subsets of cells. Morphological profiling has successfully been used to characterize genes and compounds in a number of studies. For instance, morphological profiling of chemical compounds has been used to determine their mechanism of action ^7,14–18^, identify their targets^19,20^, discover relationships with genes^20,21^, and characterize cellular heterogeneity^22^. Genes have been analyzed by creating profiles of cell populations where the gene is perturbed by RNA interference, which in turn have been used to cluster genes^23,24^, identify genetic interactions^25–27^, or characterize cellular heterogeneity^28^.

### Development of the protocol

Until recently, most published profiling methods (such as those cited above) were performed on assays involving only three dyes. We sought to devise a single assay illuminating as many biologically relevant morphological features as possible, while still maintaining compatibility with standard high-throughput microscopes. We also wanted the assay to be feasible for large-scale experiments in terms of cost and assay complexity, so we chose dyes rather than antibodies. After considerable assay development, we selected six fluorescent stains imaged in five channels, revealing eight cellular components or compartments in a single microscopy-based assay^29^ (Figure 1). We later dubbed the assay “Cell Painting”, given our aim to paint the cell as richly as possible with dyes. Automated image analysis pipelines extract ~1000-1500 morphological features from each stained and imaged cell to produce profiles. Profiles are then compared against each other and mined to address the biological question at hand. The Cell Painting assay described in this protocol has been successfully employed by multiple researchers. It was developed at the Broad Institute, carried out in multiple laboratories there, and later independently adopted at Recursion Pharmaceuticals; this protocol thus summarizes the implementation of the protocol at two independent sites and by more than ten different researchers.

**Figure 1:**
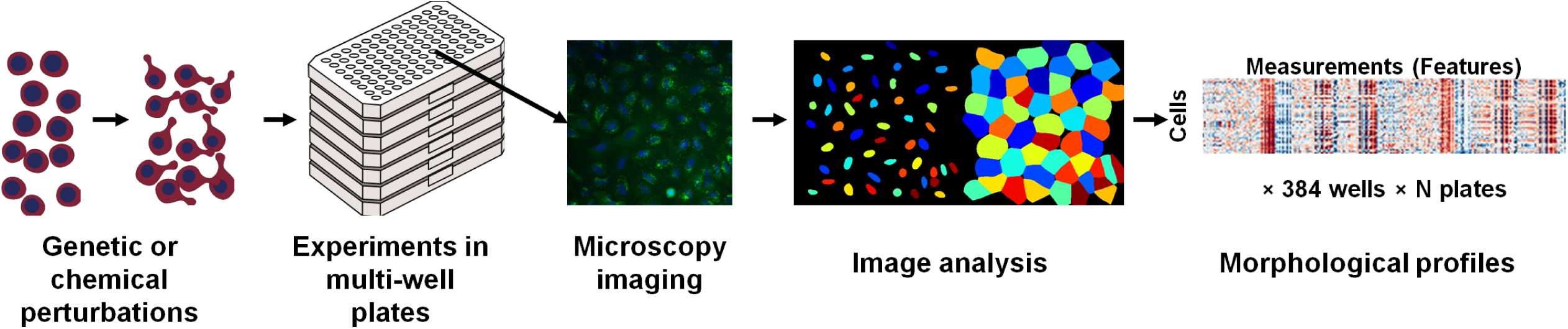
Overview of the strategy of morphological profiling using an image-based assay. After perturbing, staining, and imaging cells, the open-source software CellProfiler is used to extract 1,000+ morphological features of each cell. The collection of features is known as a *profile*: it reflects the phenotypic state of the cells in that sample, and can be compared to other profiles to make inferences.

### Applications of the method

Morphological profiling using the Cell Painting assay may be tremendously powerful for achieving a number of biological goals, only some of which have been demonstrated so far.

First, clustering small molecules by phenotypic similarity using the Cell Painting assay is effective. The first paper to use the protocol was a proof of principle study wherein cells were treated with various small molecules, stained and imaged using the Cell Painting assay, and the resulting profiles were clustered to identify which small molecules yield similar phenotypic effects^29^. Thus, the assay could be used to identify the mechanism of action or target of an unannotated compound (based on similarity to well-annotated compounds) or to “lead hop” to find additional small molecules with the same phenotypic effects but different structures (based on phenotypic similarity to compounds in a library with more favorable structural properties). As well, small molecule hits from a screen could be clustered based on morphological profiles in order to reveal potential differences among hit classes in terms of mechanism as well as polypharmacology (e.g., off-target effects).

Likewise, similarities among genetic perturbations can reveal their biological functions, by matching unannotated genes to known genes based on similar phenotypic profiles derived from the Cell Painting assay. Due to caveats concerning RNAi off-target effects^30^ (see “Limitations”), in our current work (Broad Institute), we instead overexpress genes and mine for similarities in the phenotypic profiles thus induced. In addition to mapping unannotated genes to known pathways based on profile similarity, overexpressing variant alleles is likely to enable discovery of the functional impact of a genetic variant by comparing profiles induced by wild-type and variant versions of the same gene.

Cell Painting can also be used first to identify a phenotypic signature associated with disease, then as a screen to revert that signature back to “wild-type”. Our laboratory (Recursion Pharmaceuticals) has implemented this approach in systematic fashion by simultaneously modeling hundreds of rare, monogenic loss-of-function diseases in human cells. We identify diseases for which we find a strong disease-specific phenotype in the Cell Painting assay. We then screen a drug-repurposing library to identify drugs that can reduce the strength of the disease phenotype and thus rescue the putative disease-specific features of the profile. Ultimately, the goal is to find new indications for existing drugs; this general approach (using an assay of three stains rather than the Cell Painting assay) has already been used successfully to identify potential new uses of known drugs for the treatment of cerebral cavernous malformation, a hereditary stroke syndrome^31,32^.

Lastly, both academic and pharmaceutical screening groups have an increasing desire to improve the efficiency of screening, particularly when assays are complex and expensive^33^. Profiles generated by Cell Painting applied to a large set of small molecules can be used to identify a more efficient, enriched screening set that minimizes phenotypic redundancy. The benefit is to maximize profile diversity (and thus likelihood of diverse phenotypic effects) while simultaneously eliminating compounds that do not produce any measurable effects on the cell type of interest. In a recent study, morphological profiling by Cell Painting was more powerful for this purpose than choosing a screening set based on structural diversity or diversity in high-throughput gene expression profiles^34^.

### Comparison with other methods

Though diverse methods for generating rich (100+ measurement) profiles of biological samples exist, such as metabolomic or proteomic profiling, to our knowledge gene expression profiling by L1000 (http://www.lincscloud.org/l1000/, ^35^) is currently the only practical alternative to image-based morphological profiling, in terms of throughput and efficiency^4^. Both profiling approaches (morphological profiling by Cell Painting and gene expression profiling by L1000) yield ~1000 raw features for each sample. Both are likely to capture a broad range of cellular states following perturbation. Currently, gene-expression profiling in high-throughput can be performed only on cell population aggregates and not at the single cell level, whereas morphological profiles are obtained at the level of individual cells, potentially improving the ability to resolve changes in subpopulations of cells. Cell Painting is also currently less costly per sample.

Beyond the examples above, many interesting points of comparison have yet to be rigorously tested. Thus far, no quantitative comparison of the two profiling methods’ reproducibility has been published. The two methods likely have quantitatively and qualitatively different information content; only one direct comparison has been published so far, the above-mentioned study indicating better predictive power for Cell Painting vs. L1000 gene expression profiling, for the purposes of library enrichment^34^. The study also indicated that orthogonal profiling approaches are capable of capturing a wider range of biological performance diversity than either technique alone. The fact that the two profiling methods yielded only partially overlapping library selections indicates the two modalities capture distinct information about cell state and thus are likely quite complementary. No studies have been published yet combining the two orthogonal modalities into a single profile; we believe this approach would be extremely powerful.

An alternative, image-based approach to the Cell Painting assay described here is to use a different selection of stains. The choice of stains for the Cell Painting protocol was based on the desire to detect a broad range of phenotypic effects upon compound treatment or genetic perturbation, while keeping the assay inexpensive and straightforward to implement using conventional sample preparation and imaging equipment. Morphological profiling has proven to be powerful using a relatively diverse and unbiased set of stains, that is, stains not selected to target particular pathways. Differences in phenotypic signature can be detectable even though the Cell Painting stains do not, a priori, seem likely to label a pathway targeted by a particular perturbation^14,29^. If an experiment’s questions are targeted to a particular biological area of interest, it is worth considering replacing one or more of the original stains for a more targeted one; the same basic principles of morphological profiling will still apply. However, exchanging stains entails sometimes significant optimization effort; for example, replacing the Alexa 488-ConA label requires a fluorophore bright enough to mask the fluorescence of SYTO 14 in the FITC/GFP channels (see Table 1, note 2). The Cell Painting assay could also be adapted for use with living cells, imaged over time, by using an alternative set of live-cell compatible stains or expressing fluorescently tagged proteins.

**Table 1:** Details of the channels and stains imaged in the Cell Painting assay.

| Filter<br>(excitation/emission) | Exposure<sup>1</sup><br>(ms) | Dye | Organelle or cellular<br>component | Channel<br>name, in<br>CellProfiler |
| --- | --- | --- | --- | --- |
| DAPI (387/447 nm) | 100 | Hoechst 33342 | Nucleus | DNA |
| GFP (472/520 nm)<sup>2</sup> | 100 | concanavalin A (con A)<br>AlexaFluor488 conjugate | Endoplasmic reticulum<sup>2</sup> | ER |
| Cy3 (531/593 nm) | 200 | SYTO 14 green fluorescent<br>nucleic acid stain | Nucleoli, cytoplasmic<br>RNA<sup>3</sup> | RNA |
| TexasRed (562/624<br>nm)<sup>4</sup> | 100 | Alexa Fluor 594 phalloidin<br>conjugate, wheat germ<br>agglutinin (WGA) AlexaFluor594<br>conjugate | F-actin cytoskeleton,<br>Golgi, plasma<br>membrane | AGP |
| Cy5 (628/692 nm) | 400 | MitoTracker Deep Red | Mitochondria | Mito |

^1^ The exposure times shown are those typically used by Recursion Pharmaceuticals. These values should be optimized for each instrument and/or staining procedure.

^2^ Alternately, a FITC (482/536) filter may be used.

^3^ The reagent SYTO 14 was selected after comparing a number of RNA-staining SYTO dyes because it appeared to have the highest nucleolar RNA affinity and the lowest cytoplasmic RNA affinity. Though unbound SYTO 14 fluoresces primarily in the green spectrum, the excitation/emission maxima for SYTO 14 bound to RNA is 521/547 nm, making the Cy3 channel more appropriate to use for nucleoli detection than GFP. Some cytoplasmic staining is still noticeable in the Cy3 channel and nucleolar staining is noticeable in the GFP channel.

^4^ In ^29^, the TexasRed filter was incorrectly listed as excitation/emission (562/642 nm); it is actually (562/624 nm).

### Experimental Design

#### Cell type

We and our collaborators have successfully applied the Cell Painting assay to 13 different cell cultures, including cell lines, primary cells and co-culture systems. Although we have used U2OS and A549 cells most commonly, the staining protocol has worked well in our hands for MCF-7, 3T3, HTB-9, HeLa, HepG2, HEKTE, SH-SY5Y, HUVEC, HMVEC, primary human fibroblasts, and primary human hepatocyte/3T3-J2 fibroblast co-cultures.

In many applications, the assay is intended to be unbiased and not targeted to a particular biological area of interest; in such cases using one of these already-tested cell types is sensible. In profiling, it is anticipated that many very specific biological effects and pathways can be interrogated even in a relatively “generic” cell type^13,29^.

In other cases, it is worth considering selection of a cell line that is physiologically relevant and well-characterized in the biological area of study; for example, avoiding immortal cell lines with significant genetic alterations might be essential for certain kinds of studies. It may also be valuable to choose a cell line for which there are additional sources of complementary data available, if data integration is a goal. Generally no adjustment of the staining protocol has been needed from cell type to cell type; the only change in protocol is to optimize the seeding density to adjust the confluency of the cells in accordance with the perturbations applied and biology to be examined (see “Perturbations and/or timepoints ” below).

Because we do not consider it risky to apply the assay to new cell types, selecting a new cell type might be warranted, subject to the following criteria. A major criterion in choosing a cell line is that a seeding density can be identified at which individual cells do not substantially or frequently overlap each other in the final images, that is, that the cells form a monolayer. This will allow accurate measurements to be obtained from single cells upon image analysis. It should be noted, however, that it may still be possible to obtain rich and useful data *without* the accurate segmentation of individual cells. This has been demonstrated to work for some applications^36–38^, but whether this suffices for intensive morphological profiling applications remains untested.

A second major criterion is that the cell type should grow in a manner conducive to fluorescent imaging and analysis. Specifically, the cells should typically be adherent and grow reasonably flat (i.e., non-spheroid), without significant clumping under the culture conditions used. Cell types we have tested that fail to meet this criterion are SW480 and DLD-1; presumably, non-adherent cells that are grown in suspension would also be less than ideal. The more rounded a cell type is, or the more cells grow on top of each other, the less internal structure is clearly visible by microscopy. In such cases, the staining protocol itself will label the appropriate components and images can be produced and processed, but the information content is likely to be lower for cell types with a rounded morphology as compared to a more flattened cell type.

#### Plate layout and selection of replicates and controls

When selecting the plate layout (that is, the pattern of treatments and controls across each multi-well plate), and the number of controls and replicates, the predominant concern is that phenotypic effects may be subtle. Therefore, the experiment requires careful design to avoid the impact of systematic errors^39,40^.

In order to test and compensate for systematic effects related to well position, replicates should be present in at least two different positions on a plate. This can be accomplished by having either two (or more) replicates on the same plate, or having two (or more) plate layouts where the same set of perturbations are present in different well positions. Sample replicates should not be placed adjacent to each other on the plate (especially controls) but instead spaced out as much as possible. Having all the replicates of a perturbation (or a control) lying on the edge of the plate should also be avoided.

At least four replicates are recommended as we (Broad Institute) have observed a significant loss in data quality with fewer replicates. In practice, we have used five or more replicates to buffer against accidental sample loss. For larger scale experiments (1000+ perturbations), we have used four replicates for cost reasons.

Control wells should be scattered across the plate rather than having them clustered in neighboring wells or placed in a single row or column. The standard pattern used at the Broad Institute for compound screening is to place the control wells in a chevron pattern across the plate; other patterning variations are feasible. Ideally, some control wells should be placed in a pattern that can be distinguished if the plate is turned 180°, e.g., placing a particular cytotoxic treatment in the top left well but not in the bottom right well, or omitting cells at several non-symmetric well locations.

For morphological profiling, negative controls are used to normalize the image features, but positive controls can be included if they can be reliably defined for the experiment. For compound library screening, typically the negative controls are ‘vehicle’-only conditions (e.g., DMSO). We (Broad Institute) have found that ~30 wells/plate designated for negative controls works well for a 384-well format assay. In cases where there is no obvious negative control, the untreated wells (i.e. wells containing cells but not subjected to drug vehicles nor gene perturbant delivery reagents) may serve as a substitute for this purpose. If more than one type of control is used, the positions should be interleaved. For gene profiling, multiple negative controls may be considered such as empty vectors, or control treatments towards genes that are irrelevant or non-native to the cell type. However, it should be kept in mind that in the case of gene knockdown using RNAi, even control hairpins targeting the same gene but containing different seed sequences can induce different morphological profiles, a result verified by the Cell Painting assay^30^.

With automated plate handlers, groups of plates are typically washed and stained as a single batch. We recommend not processing replicate plates within a single batch, but rather spreading them out across multiple batches. Additionally, a given replicate plate should be processed in a different order for each batch. The rationale is to keep the systemic batch effects orthogonal to any biological effects of interest that may be present.

#### Perturbations and/or timepoints

Prior to carrying out a large-scale experiment, consider initially performing a smaller pilot with a small number of perturbations/timepoints. During assay development, we recommend culturing the cells until the desired timepoint, and then examining them for the degree of confluency. This assessment may be performed by fluorescent staining (i.e., following some or all of the regular protocol) or even by eye under brightfield; the latter is less effort and is feasible once a researcher has some experience with the assay. Of note, the pilot assay should replicate the conditions under which the cells will be analyzed in the full experiment as closely as possible; at a minimum including vehicle controls for small molecule treatment and transfection or infection with scrambled control (or similar) for RNAi screens, as such perturbations can have a significant effect on cellular growth and confluence. We have used 24-hour (Recursion Pharmaceuticals) and 48-hour (Broad Institute^29,34^) small molecule exposures prior to fixation and imaging. For infection/transfection of RNAi and over-expression plasmids, we use 96 hours ^30^ and 72 hours, respectively. It may well be that a shorter or longer exposure time is optimal, particularly for certain types of biological processes of interest and certain perturbation types.

As noted above, some adjustment of the cell seeding density may be needed depending on the cell line used and the biological processes under examination. Since cell-cell junction interactions play a significant physiological role in endothelial and epithelial cell types, we (Recursion Pharmaceuticals) recommend growing these cultures as confluent or near-confluent monolayers. If such biological processes are not of particular concern, we (Broad Institute) recommend optimizing the cell density while striking a balance between two considerations. Because the expected phenotypes are subtle, a low cell count will lead to a small sample size that is not truly representative of the phenotype. In cases where a fair number of perturbations may be cytotoxic, increased seeding density may mitigate the smaller numbers of surviving cells comprising the morphological sample. On the other hand, if the cell number is too high (or the cells form a confluent monolayer in the extreme case), the cells are too crowded for representative measurements of many phenotypes, particularly for image features derived from cell shape. Therefore, we at the Broad Institute aim for a seeding density that provides the cells enough space to exhibit their full-fledged morphological phenotypes while maintaining a high sample size for each phenotype expressed. Generally, we have found that ~80% confluency at the time of fixation provides a good balance.

The assay is theoretically amenable to evaluating any biological perturbation type. We have performed the Cell Painting assay with small molecule treatments^29,34^, viral infection^30^, transient transfection, and using selectable markers resulting in all surviving cells receiving the treatment.

### Level of expertise needed to implement the protocol

Experience with high-throughput automated equipment is required to carry out the full sample preparation portion of the protocol, although in certain cases the assay might be carried out at a smaller scale manually and using a non-automated microscope. Core facilities at many institutions would likely be well-equipped to aid laboratories in conducting the Cell Painting assay at larger scale. Prior image analysis experience is helpful but not necessary to carry out the image analysis procedure on a desktop computer. However, this solution is not suitable for large-scale Cell Painting assays with greater than ~1000 images. In such cases, we recommend using a computing cluster, which will likely require an information technology (IT) expert’s assistance. An active moderated forum (http://www.cellprofiler.org/forum/) exists for answering questions and troubleshooting issues that may arise using CellProfiler. Thus, despite some technical challenges, this protocol is accessible to nearly any laboratory which, at the least, has access to collaborators with some experience with high-throughput automated screening, some advanced computational skills, and possesses a willingness to learn.

### Limitations

Although the Cell Painting assay is intended to be unbiased with regard to the cell type chosen, certain biological processes may simply not yield any relevant discernible morphological phenotypes, given the experimental conditions used (stains, cell type, time point, etc.). In this case, augmenting the image-based profiles with additional or orthogonal assays may reveal additional biological effects that would be otherwise missed. In addition, alternate stains may be chosen to highlight the relevant cellular subcompartments while maintaining broad coverage of other organelles (see “Comparison with other methods” above for more details).

For RNAi experiments, the magnitude and prevalence of off-target effects in mammalian cells via the RNAi seed-based mechanism make the complete morphological profiles of RNAi reagents targeting the same gene rarely look any more similar than those targeting different genes(^30^, ^30^related to high-dimensional profiling in general, not specifically to morphological profiling. This impedes large-scale experiments using short RNAi reagents where the experimental design requires widespread comparisons across all samples. However, we note that it does not preclude experiments where the goal is to identify particular genes for which multiple RNAi reagents do yield a consistent profile, as is the case for our work identifying disease-associated phenotypes at Recursion Pharmaceuticals. An alternative gene suppression technique, CRISPR/Cas9, has not yet been used in conjunction with morphological profiling and will likely be effective.

Finally, there are some computational challenges associated with this assay. First, there are statistical challenges associated with the analyzing the high-dimensional feature space that results from Cell Painting. Similar to the case with gene expression data^41^, issues such as the curse of dimensionality, model overfitting, spurious correlations, and multiple testing complicate the data analysis of this assay; these types of challenges are widely recognized in systems biology^42^. Second, while in principle single-cell data is preferable over aggregated data, the former requires substantially more computational storage and processing resources and furthermore, no routine analytical protocol has yet been established for this. Lastly, data analysis across separately-performed experiments is likely to be complicated, requiring proper control for the potentially substantial effects of differences in cell seeding, growth, and other batch-related or other systematic artifacts. Protocols for such cases have not yet been developed.

## MATERIALS

### REAGENTS

Note: The procedure is described as implemented in two particular laboratories. Materials listed are those used by Recursion Pharmaceuticals for brevity. Where multiple options are noted in this section, those used at the Broad Institute are indicated with an asterisk (*) and those used at Recursion Pharmaceuticals are indicated with a plus (+).

#### Cell culture

- Cells, e.g., U2OS cells (ATCC, cat. no. HTB□96) or A549 cells (ATCC, cat. no. CCL-185)
- DMEM (Fisher Scientific, cat. no. MT-10-017-CV)
- Fetal bovine serum (FBS) (Life Technologies, cat. no. 10437-028)
- Penicillin□streptomycin (Fisher Scientific, cat. no. MT-30-002-CI)
- Trypsin, TrypLE™ Express Enzyme (Life Technologies, cat. no. 12605-036)
- Phosphate Buffered Saline pH 7.4 (PBS) (Life Technologies, cat. no. 10010-023)
- Microplates: Corning 384-well black/clear flat bottom, fibronectin-coated (Corning, cat. no. 4585)^+^, Corning 384-well black/clear flat bottom, TC-treated, bar-coded (Corning, cat. no. 3712BC)*

#### (Optional) siRNA transfection

- Lipofectamine RNAiMax (Life Technologies cat. no. 13778030)
- Silencer Select Pre-designed and custom siRNAs (Ambion) **Caution:** The above reagents work well for many cell types, but lengthy optimization is often required in new cell types for which siRNA transfection is performed. Specific conditions of transfection should be evaluated in pilot assays to confirm suitability.
- Optimem (Life Technologies, cat. no. 31985-070)
- Deep 384-well plates (USA Scientific, cat. no. 1884-2410)

#### (Optional) Compound treatment

- Small-molecule libraries, typically 10 mM stock in DMSO (e.g., Chembridge Library or Maybridge Library) **Caution:** Some small-molecule libraries may contain toxic compounds; suitable precautions should be taken.

#### Fixation and staining

- MitoTracker Deep Red (Invitrogen, cat. no. M22426) **Caution:** The MitoTracker stock solution is in DMSO. DMSO is a toxic chemical and easily penetrates the skin. One must avoid ingestion, inhalation and direct contact with skin and eyes. Use proper gloves to handle DMSO. Follow your institutional guidelines for using and discarding waste chemicals.
- Wheat germ agglutinin Alexa 594 conjugate (Invitrogen, cat.no. W11262)
- Paraformaldehyde 16%, methanol free (Electron Microscopy Sciences, cat. no. 15710□S) **Caution:** PFA is a very toxic chemical and one must avoid inhalation and/or direct contact with skin and eyes. Use proper gloves and a mask to handle PFA. Follow your institutional guidelines for using and discarding waste chemicals.
- Hank’s Balanced Salt Solution (10x), HBSS (Invitrogen, cat. no. 14065□056)
- Triton X-100 (Sigma, cat. no. T8787) **Caution:** Triton X-100 is a toxic chemical and one must avoid inhalation and/or direct contact with skin and eyes. Use proper gloves to handle Triton X-100. Follow your institutional guidelines for using and discarding waste chemicals.
- Phalloidin Alexa594 conjugate (Invitrogen, cat. no. A12381)
- Concavalin A Alexa488 conjugate (Invitrogen, cat. no. C11252)
- Hoechst 33342 (Invitrogen, cat. no. H3570)
- SYTO 14 green fluorescent nucleic acid stain (Invitrogen, cat.no. S7576)
- Sodium biocarbonate (HyClone, cat. no. SH30033.01)
- Methanol (VMR, cat. no. BDH1135) **Caution:** Methanol is a very toxic chemical and one must avoid ingestion, inhalation and/or direct contact with skin and eyes. Use proper gloves and a mask to handle methanol. Follow your institutional guidelines for using and discarding waste chemicals.
- BSA (Equitech-Bio, cat. no. BAH66)
- Aluminum single tab foil, standard size (USA Scientific, cat. no. 2938-4100)

### EQUIPMENT

- Tissue culture incubator at 37 °C, 5% CO_2_
- CyBi-Well 96/384-channel simultaneous pipettor (CyBio, cat. no. 3391 3 4112)
- Automated liquid handler: Multidrop Combi reagent dispenser (Thermo Scientific, cat. no. 5840300)*, Freedom EVO with 384-channel arm (Tecan, cat no. MCA384)^+^
- Plate washer: Biotek ELx405 HT*
- Centrifuge: Allegra 6 (Beckman Coulter, cat. no. 366802)*, PlateFuge (Benchmark Scientific, cat. no. C2000)^+^
- ImageXpress Micro XLS epifluorescent microscope (Molecular Devices)
- CRS CataLyst Express robot microplate handler system (Thermo Scientific)
- Microscope light source: LED light engine (Lumencor)
- Access to a high-performance computer or a remote-host computing cluster (optional; recommended if planning to acquire >1000 fields of view)
- CellProfiler and CellProfiler Analyst biological image analysis software; see “Equipment setup” for how to obtain
- CellProfiler pipelines; see “Equipment setup” for how to obtain

### REAGENT SETUP

#### Cell culture

- If using a cell line for the first time, we suggest testing the staining protocol in a few pilot plates in order to visually confirm visibility of the cellular features.

#### MitoTracker Deep Red stock

- The product from Invitrogen (cat. no. M22426) contains 50 μg in each vial. Add 91 μL DMSO to one vial to make 1 mM solution. Store the solution at ≤ −20°C, protected from light, and use it within one month.

#### Wheat Germ Agglutinin (WGA) Alexa594 conjugate stock

- The product from Invitrogen (cat. no. W11262) contains 5 mg in each vial. Add 5 ml dH2O to each to make 1 mg/ml solution. Store the solution at −20°C, protected from light, and use it within one month. We recommend centrifuging the WGA conjugate solution briefly before use, in order to remove any protein aggregates in solution which would contribute to nonspecific background staining.
- We (Recursion Pharmaceuticals) note that using the concentrations in the Gustafsdottir paper^29^, the actin fibers labeled by the phalloidin staining were not discernible in the TexasRed channel due to excessive brightness of the WGA Golgi/plasma membrane staining. The stated reagent concentrations suffice as a starting point, but required adjustment in our laboratory, resulting in a WGA concentration of 5 μg/mL for live cell staining and 0.1-0.5 μg/mL for fixed and permeabilized cells to achieve the desired effect.

#### Concanavalin A Alexa488 conjugate stock

- The product from Invitrogen (cat. no. C11252) contains 5 mg in each vial. Add 5 ml 0.1 M sodium bicarbonate to each vial to make 1 mg/ml solution. Store the solution at −20°C, protected from light, and use it within one month.

#### Phalloidin Alexa594 conjugate stock

- The product from Invitrogen (cat. no. A12381) contains 300 units in each vial. Add 1.5 ml methanol to each vial. Store the solution at −20°C, protected from light, and use it within one year.

#### SYTO 14 green fluorescent nucleic acid stain stock

- The product from Invitrogen (cat.no. S7576) is 5 mM solution in DMSO. Store the solution at −20°C, protected from light, and use it within one year.

#### Hoechst 33342 stock

- The product from Invitrogen (cat. no. H3570) is 10 mg/ml solution in water. Store the solution at 4°C, protected from light, and use within six months.

#### HBSS (1X)

- The product from Invitrogen (cat. no. 14065-056) is 10X. Add 100mL HBSS (1X) to 900mL water to make HBSS (1X). Filter the HBSS (1X) with 0.22 μm filter.

#### 1% BSA solution in HBSS

- Weigh 1g BSA and dissolve in 100 ml HBSS to make 1% (wt/vol) BSA solution. Filter the 1% BSA solution with a 0.22 μm filter. Make fresh solution for each experiment.

#### 0.1% Triton X-100 solution in HBSS

- Add 100uL Triton X-100 into 100 ml HBSS to make 0.1% (vol/vol) Triton X-100 solution. Make fresh solution for each experiment.

#### Compound library

- Dissolve the compounds in DMSO to yield the desired molarity; the final concentration should be such that the density is equivalent to the cell culture media. Seal and store at −20 °C for long-term storage or at RT for up to 6 months.

### EQUIPMENT SETUP

#### Microscope selection

- We use an ImageXpress Micro XLS epifluorescent microscope (Molecular Devices). Images are captured in 5 fluorescent channels (ImageXpress filters are listed, followed by excitation/emission in nm): DAPI (387/447), GFP (472/520) (or alternately, FITC (482/536)), Cy3 (531/593), TexasRed (562/624), Cy5 (628/692). A sample ImageXpress plate acquisition settings file is available as Supplementary Data 1. **Critical:** The same microscope should be used for imaging all microtiter plates during an experiment. We do not recommend switching microscopes mid-stream for a batch of images to be analyzed because lamp intensities, filter patterns and other subtleties can be quite different even between supposedly identical microscope setups.
- If multiple microscopes must be used, we recommend imaging one full replicate all on one microscope, as opposed to arbitrarily assigning plates to different instruments as the experiment proceeds. The rationale is to avoid imager-induced batch effects. If the differences between perturbations are dramatic, then post-acquisition normalization will probably be effective (see Step 5, “Normalize morphological features across plates” for more details). However, if the morphological effects to be measured are subtle, normalization may not be sufficient, and the similarities in the collected image features will more likely reflect the different image acquisition than the underlying biological perturbations.

#### Automated image acquisition settings

- The images should be acquired with the maximum bit-depth possible, in a “lossless” image format so as to preserve all the information captured by the light sensor. Each channel should be captured as an individual grayscale image. For the ImageXpress XLS system, a 16-bit grayscale TIFF is sufficient. No further preprocessing should be performed on the images prior to analysis.
- Typically, nine sites are collected per well in a 3 x 3 site layout, at 20x magnification. Time permitting, more sites can be imaged in order to increase well coverage and improve sample statistics; it is best to capture as many cells as possible. **Critical:** Avoid capturing the edges of the well in the images, particularly if a large number of sites/well are imaged. While it is feasible to remove the well edges from the images post-acquisition using image processing approaches, such methods are challenging and best avoided. One helpful approach is to reduce the field of view size in order to avoid the well edges; this setting (expressed as a percentage) is accessible through the “Sites to Visit” tab in the MetaXpress software.
- If using an ImageXpress microscope, use laser-based focusing with image recovery for autofocusing. The autofocusing can be applied for only the first site of each well, or alternately for all sites; the latter is a minimal increase in time and is recommended for those using glass-bottom plates to decrease focussing problems.
- Use the Hoechst channel (DAPI, “w1” in ImageXpress) for the image recovery, with the focus binning set to 3 and a Z-offset for the other channels. The choice of optimal Z-offset will depend on the cell line and should be set by looking for the optimal focus by eye.
- Suggested exposure times (Recursion Pharmaceuticals) are given in the Table 1. However, these values should be optimized for each experiment. Higher exposure times will yield a larger dynamic range but will increase automated image acquisition time. **Critical:** Be sure that the images are not saturated. Generally, set exposure times such that a typical image uses roughly 50% of the dynamic range. For example, because the pixel intensities will range from 0 to 65,535 for a 16-bit image, a rule-of-thumb is for the typical sample to yield a maximum intensity of ~32,000. This guideline will prevent saturation (i.e., hitting the value 65,535) from samples that are brighter due to a perturbation.
- Do not use shading correction, as the background illumination heterogeneities will be corrected post-acquisition using the CellProfiler software.
- Prior to beginning the complete imaging run, it is useful to capture images from 3 to 5 wells at a few different locations across the plate, in order to confirm that the microscope is operating as expected and the acquisition settings are optimal for the experiment and cell line at hand.
- We recommend exporting the microscope configuration for future use once the optimal settings have been determined. A sample ImageXpress plate acquisition settings file is provided as Supplementary Data 1.
- An example image data set for an RNAi Cell Painting study of U2OS cells is supplied in Supplementary Note 1.

#### Image processing software

- CellProfiler biological image analysis software is used to extract per-cell morphology feature data from the Cell Painting images, as well as per-image quality control metrics.
- To download and install the open-source CellProfiler software, go to http://cellprofiler.org, follow the download links for CellProfiler and follow the installation instructions. The current version at the time of writing is 2.1.1.
- This protocol assumes basic knowledge of the CellProfiler image analysis software package. Extensive online documentation and tutorials can be found at http://www.cellprofiler.org/. Also, the ‘?’ buttons within CellProfiler’s interface provide detailed help. The pipelines used here are compatible with CellProfiler version 2.1.1 and above.
- This protocol uses three CellProfiler pipelines to perform the following tasks: (a) illumination correction, (b) quality control; and (c) morphological feature extraction. These are available as Supplementary Data 2 and continuously updated versions are freely available for download at http://cellprofiler.org/examples.shtml.
- The pipelines assume that image files follow the nomenclature of the ImageXpress microscope system, in which the plate/well/site metadata is encoded as part of the filename. The plate and well metadata in particular are essential because CellProfiler uses the plate metadata in order to process the images on a per-plate basis, and the plate and well metadata are needed for linking the plate layout information with the images for the downstream profiling analysis. Therefore, images coming from a different acquisition system may require adjustments to the Metadata module to capture this information; please refer to the help for this module for more details.
- The quality control and morphological feature extraction pipelines are set to write out cellular features to a MySQL database which is recommended for a analyses involved > 1000 images; see “Computing system” for details. If using a smaller number of images, the pipelines can be adjusted to output the measured features to a comma-delimited file (CSV) using the *ExportToSpreadsheet* module. Third-party data analysis tools may be more amenable to importing data from a CSV-formatted file than from a MySQL database. However, the scripts provided in Supplementary Methods 2 to generate per-well profiles from the extracted features are MySQL-only.

#### Computing system

- If the number of images to analyze is sufficiently large that a single computer would take too long to process them (more than ~1000), we recommend using a computing cluster if available, e.g, a high-performance server farm or a cloud-computing platform such as Amazon AWS. Carrying out this step requires significant setup effort and will probably require enlisting the help of your IT department. Please refer to the Supplementary Methods 1 or to our GitHub webpage https://github.com/CellProfiler/CellProfiler/wiki/Adapting-CellProfiler-to-a-LIMS-environment for more details.
- We recommend setting up a MySQL database to allow multiple CellProfiler processes to write out cellular features in parallel; doing so will probably require enlisting the help of your IT department. This database will be used by the CellProfiler module *ExportToDatabase* to create data tables as described under Procedures, as well as by the scripts to generate per-well profiles from the extracted features (see Supplementary Methods 2).

#### Image data exploration software

- The CellProfiler Analyst data exploration software may be used to explore the data or for quality control^43^.
- To download and install the CellProfiler Analyst software, go to http://cellprofiler.org, follow the download link for CellProfiler Analyst and follow the installation instructions. The current version of CellProfiler Analyst at the time of writing is 2.0.

## PROCEDURE

### 1. Cell Culture

**Timing: variable; 4 - 7 d**

a. The following cell-plating procedure is validated for many cell types; each step may need adjustment depending on local conditions or alternate cell types. We have included recommended optional steps for experiments involving siRNA transfection.
b. Prepare cells for seeding according to known best practices for the cell type of choice. The following protocol is validated for use on A549 cells in Corning 384-well 200nM thick glass bottomed plates.
c. Grow cells to near confluence (~80%) in a T-150 culture vessel.
d. Optional: Prepare siRNA transfection. The following protocol is written for a 10 nM final siRNA concentration in 0.1% Lipofectamine RNAiMAX which achieves >70% knockdown for diverse targets in A549, U2OS, and HUVEC cells.
  i. Thaw and dilute siRNAs to a concentration of 2 μM in sterile molecular-grade water.
  ii. Dilute Lipofectamine RNAiMAX with FBS-free Opti-MEM medium (1:100) in appropriate RNase-free deep 384-well plates.
  iii. Mix 0.2 μL diluted siRNA with 10 μL diluted Lipofectamine RNAiMAX.
  iv. Ensure that siRNA-Lipofectamine mix has incubated at room temperature for at least 10min before proceeding to step i.
e. Rinse cells with PBS without Ca^2+^or Mg^2+^.
f. Add 6 ml TrypLE Express and incubate at room temperature for 30 seconds. Remove the TrypLE Express and add 1 ml fresh TrypLE Express. Incubate at 37°C until the cells have detached. This should occur within 3-5 minutes.
g. Add 10 ml growth medium to deactivate trypsin, and determine the live cell concentration using standard methods (hemocytometer or cell counter).
h. Dilute A549 cells to 50,000 live cells/ml in media, and dispense 40 μl (2000 live A549 cells) to each well of the 384-well plates using a Multidrop Combi reagent dispenser. Different cell types and growth conditions will require variations in seeding density; typical ranges will vary from 1500 to 3000 cells/well. **Critical step:** Adequately resuspend the cell mixture to ensure a homogeneous cell suspension prior to each dispense. It is not uncommon for cells to rapidly settle in their reservoir resulting in plate-to-plate variation in cell numbers. If utilizing a liquid handler with a multi-dispense function, be sure to adequately prime the dispensing cassette and/or dispense at least 10 μL of cell suspension back into the reservoir prior to dispensing the cells into culture plates; the latter is helpful if cells or reagent are sticking to the tubing. **Critical step:** When handling liquid for many plates with one set of tips, confirm that no residual bubbles within the tips touch the head of the liquid handler during aspiration in order to ensure accurate liquid dispensation.
i. Optional: Add the siRNA-transfection reagent mixture to the 384-well plate using the CyBi-Well simultaneous pipettor, at 10.2 μL /well.
j. Allow plates to set on a flat, level surface at room temperature for 1 - 2 hrs to reduce plate edge effects^44^.
k. Put the plates into the incubator (37°C, 5% CO_2_, 90-95% humidity). To reduce plate edge effects produced by incubator temperature variations and media evaporation, we recommend either spacing out the plates in the incubator or using racks with “dummy” plates filled with liquid placed on the top and bottom. We also recommend rotating the plates/stacks within the incubator to avoid positional effects.
l. Replace the culture medium with 50 μl of 2% FBS in DMEM 24 hours after seeding. Perform the aspiration steps using a plate washer such as the BioTek ELx405 microplate washer or equivalent. Reducing FBS concentration minimizes the risk of overgrowth at the time of fixation. Optimal FBS concentrations may vary depend on the cell type and transfection reagent selected.
m. Optional: Add starvation medium (0.01% FBS in DMEM) to the cells approximately 24 hours prior to staining and fixation in order to synchronize cell growth rate. If siRNA transfection is included in previous steps, we (Recursion) have found that addition of starvation medium 72 hours after transfection results in robust knock-down in a number of cell types. Cell cycle synchronization may improve profile quality because it reduces variability in whole-culture cell cycle stage; however, profiling of asynchronous cell populations may enable capture of phenotypes affecting all stages of the cell cycle.
n. Optional: Add compounds to cells using a pin tool or liquid handler. We add compounds 24 hours (Recursion Pharmaceuticals) or 48 hours (Broad Institute) prior to staining and fixation, but the timing should be adjusted depending on the growth rate of each cell type and the biological processes under consideration. Recursion Pharmaceuticals typically adds compounds to cells in an environment that is antibiotic-free (to avoid perturbations arising from complex antibiotic-drug interactions) and low-serum (to synchronize cell state). To ensure adequate mixture of compounds in solution, we recommend that compounds are mixed well in the culture medium before adding to cells.

### 2. Staining and Fixation

**Timing: variable; 2.5-3 h for one batch experiment of 384-well plates**.

a. Live cell MitoTracker and Wheat Germ Agglutinin staining.
  i. Prepare a 500 nM MitoTracker, 60 μg/ml WGA solution in prewarmed media (DMEM, 10% FBS). To make 50 ml MitoTracker/WGA staining solution, add 19.3 μL MitoTracker stock solution and 2.275 ml WGA stock solution to 47.7 ml prewarmed media. The reason for applying the WGA solution at the live cell stage is that we (Broad Institute) found that if the cells are stained with WGA after fixation, the WGA staining pattern becomes distorted after the plates are stored in refrigeration, even after a couple of days at 4°C; live cell staining fixed this problem.
  ii. Remove media from plates, set the aspiration height in the plate washer to leave 10uL of residual volume to minimize the disturbance to the live cells from the pins and media turbulence.
  iii. Add 30 μL of MitoTracker/WGA staining solution.
  iv. Centrifuge the plate (500 g at room temperature for 1 min) after adding stain solutions and ensure there are no bubbles in the bottom of the wells.
  v. Incubate the plates for 30 min in the dark at 37 °C. **Critical Step:** Prepare the working stain solution before use. Do not store the working stain solution exposed to light or over long periods, to maintain fluorescence.
b. Fixation
  i. Add 10 μL of 16% methanol-free paraformaldehyde for a final concentration of 3.2% (vol/vol). **Critical Step:** We (Broad Institute) recommend performing the fixation and subsequent permeabilization and staining steps with no pauses. In our hands, halting between steps, e.g., between the fixing/permeabilizing steps and the staining step, results in degradation of the SYTO 14 staining quality. **Critical Step:** Having FBS or BSA present during fixation may help prevent cellular retraction.
  ii. Centrifuge the plate (500 g at room temperature for 1 min) after adding stain solutions and ensure there are no bubbles in the bottom of the wells.
  iii. Incubate the plates in the dark at room temperature for 20 min.
  iv. Wash the plates once with 70 μL 1X HBSS.
c. Permeabilization
  i. Remove HBSS and add 30 μL of 0.1% (vol/vol) Triton X-100 solution to the wells.
  ii. Centrifuge the plate (500 g at room temperature for 1 min) after adding stain solutions and ensure there are no bubbles in the bottom of the wells.
  iii. Incubate the plates in the dark at room temperature for 10-20 min.
  iv. Wash the wells twice with 70 μL 1x HBSS.
d. Phalloidin, ConcanavalinA, Hoechst, and SYTO 14 staining.
  i. Prepare a 25 μL/ml phalloidin solution, 100 μg/ml ConcanavalinA, 5 μg/ml Hoechst 33342 and 3 μM SYTO 14 green fluorescent nucleic acid stain solution in 1x HBSS, 1% BSA. To make 50 ml stain solution, add 1.25 ml phalloidin stock solution, 1 ml ConcanvalinA stock solution, 25 μL Hoechst stock solution and 30 μL SYTO 14 green fluorescent nucleic acid stain stock solution to 47.7 ml 1% BSA solution in HBSS.
  ii. Remove HBSS and add 30 μL of staining solution to each well.
  iii. Centrifuge the plate (500 g at room temperature for 1 min) after adding stain solutions and ensure there are no bubbles in the bottom of the wells.
  iv. Incubate the plates in the dark at room temperature for 30 min.
  v. Wash cells three times with 70 μL 1x HBSS, with no final aspiration.
  vi. Seal plates with adhesive foil and store at 4°C in the dark.

### 3. Automated image acquisition

**Timing: variable; ~3.5 hours per 384-well plate**

a. Mount the microtiter plates into the automated microscopy system for imaging. For large-scale Cell Painting assays, we recommend the use of an automated microplate handling system.
b. Set up the microscope acquisition settings as described in “Equipment Setup”.
c. Start the automated imaging sequence according to the microscope manufacturer’s instructions. **(TROUBLESHOOTING)**

### 4. Morphological image feature extraction from microscopy data

**Timing: variable; 20 h per batch of 384-well plates**

a. Illumination correction to improve fluorescence intensity measurements
  i. Start CellProfiler.
  ii. Load the illumination correction pipeline into CellProfiler by selecting *File > Import > Pipeline from File* from the CellProfiler main menu and selecting *illumination.cppipe*. **Critical step:** Non-homogeneous illumination introduced by microscopy optics can result in errors in cellular feature identification and can degrade the accuracy of intensity-based measurements. This is an especially important problem in light of the subtle phenotypic signatures that morphological profiling aims to capture. Non-homogeneous illumination can occur even when fiber-optic light sources are used and even if the automated microscope is set up to perform illumination correction. The use of a uniformly fluorescent reference image (“white-referencing”), while common, is not suitable to HTS. A retrospective method to correct all acquired images on a per-channel, per-plate basis is therefore recommended^45^; the illumination pipeline takes this approach.
  iii. Select the *Images* input module in the ‘Input modules’ panel to the top-left of the interface. From your file browser, drag and drop the folder(s) containing your images into the ‘File list’ panel.
  iv. Click the ‘View output settings’ button at the bottom-left of the interface. In the settings panels, select an appropriate ‘Default output folder’ where the illumination correction images will be saved.
  v. Save the current settings to a project (.cpproj) file containing the pipeline, the list of input images and the output location by selecting *File > Save Project*. Enter the desired project filename in the dialog box that appears.
  vi. Press the ‘Analyze Images’ button at the bottom-left of the interface. A progress bar in the bottom-right will indicate the estimated time of completion. **Critical step:** This step assumes that you will be running the illumination correction pipeline locally on your computer. If your institution has a shared high-performance computing cluster, we recommend executing the pipeline on the cluster as a batch process, i.e., a series of smaller processes entered at the command line; this will result in much more efficient processing. Enlist the help of your institution’s IT department to find out whether this is an option and what resources are available. If so, carry out optional Step (d), describing modifications to the pipeline to run it as a batch process.
  vii. The end result of this step will be a collection of illumination correction images in the Default output folder, one for each plate and channel. An example set of illumination correction images is provided as Supplementary Data 3.
b. Quality control to identify and exclude aberrant images
  i. Start CellProfiler, if not already running.
  ii. Load the quality control pipeline into CellProfiler by selecting *File > Import > Pipeline from File* from the CellProfiler main menu and selecting *qc.cppipe*. **Critical step:** As mentioned above, high-quality images are essential for robust downstream analysis of Cell Painting data. Therefore, we recommend implementing quality control (QC) measures. The approach detailed here uses CellProfiler to analyze the data using quality control metrics that do not require cell identification (a.k.a. segmentation in the analysis pipeline, step 4c) ^43^. However, the same goal can be met with other analytical approaches after cell identification and measurement in step 4c.
  iii. Select the *Images* input module in the ‘Input modules’ panel to the top-left of the interface. From your file browser, drag and drop the folder(s) containing your images into the ‘File list’ panel (a shortcut for this step is to simply use the same project file as the illumination step above and load the QC pipeline to replace the illumination pipeline, while retaining the same image list).
  iv. Select the *ExportToDatabase* module. The setting for “Database name” is highlighted in red because it is waiting for a proper value to be provided. Change the fields for “Database name”, “Database host”, “Username” and “Password” to their respective values appropriate for your MySQL database server; the red text will disappear at that point. Once done, you can press the “Test connection” button to confirm that the settings are correct. The setting “Table Prefix” should be changed to a different value such that a new table is created and used for this experiment.
  v. Click the ‘View output settings’ button at the top-left of the interface. In the settings panels, select an appropriate ‘Default output folder’ where the QC data will be saved.
  vi. Optional: For smaller number of images (e.g., < 1000) or instances when a MySQL database is not available, the *ExportToSpreadsheet* module may be used to export the collected measurements to a comma-delimited file (CSV) format. Add this module by selecting *Edit > Add Module > File Processing > ExportToSpreadsheet* from the CellProfiler main menu, and position it as the last module in the pipeline by using the “^” or “v” buttons beneath the pipeline. Disable the *ExportToDatabase* module by clicking the green checkmark next to the module name; the green checkmark will then be grayed out to indicate its status. Select the *ExportToSpreadsheet* module, select “No” for “Export all Measurement types?”, and then select “Image” from the “Data to export” drop-down drop that appears. This will place a CSV file containing the per-image data in the Default output folder.
  vii. Save the current settings to a project (.cpproj) file containing the pipeline, the list of input images and the output location by selecting *File > Save Project*. Enter the desired project filename in the dialog box that appears.
  viii. Press the ‘Analyze Images’ button at the bottom-left of the interface. A progress bar in the bottom-left will indicate the estimated time of completion. **Critical step:** This step assumes that you will be running the quality control pipeline locally on your computer. If your institution has a shared high-performance computing cluster, we recommend executing the pipeline on the cluster as a batch process, i.e., a series of smaller processes entered at the command line; this will result in much more efficient processing. Enlist the help of your institution’s IT department to find out whether this is an option and what resources are available. If so, carry out optional Step (d), describing modifications to the pipeline to run it as a batch process.
  ix. When the QC processing run is completed, apply the workflow described in ^43^ to use CellProfiler Analyst to explore the data and select QC image features and thresholds in order to exclude out-of-focus and saturated images from further analysis.
c. Image analysis to extract morphological features
  i. Start CellProfiler, if not already running.
  ii. Load the analysis pipeline into CellProfiler by selecting *File > Import > Pipeline from File* from the CellProfiler main menu and selecting *analysis.cppipe*.
  iii. Select the *Images* input module in the ‘Input modules’ panel to the top-left of the interface. From your file browser, drag and drop the folder(s) containing your images into the ‘File list’ panel (A shortcut for this step is to simply use the same project file as the illumination step above and load the analysis pipeline to replace the illumination pipeline while retaining the same image list). For this step, you should also drag and drop the folder containing your illumination correction images into the ‘File list’ panel.
  iv. Select the *FlagImages* module, which is used to tag images with a metadata label containing a 1 or 0, depending on whether features from particular image channels pass or fail chosen QC criteria respectively. Two sample measurements and thresholds are provided in the pipeline; the choice of QC image feature(s) and threshold(s) should be adjusted to reflect your results from step (b). To add more QC image features to an existing flag, press the “Add another measurement” button, and select “ImageQuality” as the category, the desired measurement and image from the respective drop-box boxes, whether the image is flagged based on a high or low threshold values, and the actual threshold value for the measurement. You can also add more flags by pressing the “Add another flag” button, give the flag a name and specify whether the image needs to pass any or all of the criteria to be flagged; you may add as many flags and/or features to a flag as needed. If you do not wish to use this module for QC, you can disable the module by clicking the green checkmark to the left of the module name; the checkmark is grayed out when the module is disabled.
  v. Select the *ExportToDatabase* module, which is used to write image-based feature measurements to a MySQL database. The setting for “Database name” is highlighted in red because it is waiting for a proper value to be provided. Change the fields for “Database name”, “Database host”, “Username” and “Password” to their respective values appropriate for your MySQL database server; the red text will disappear at that point. For provenance purposes, we recommend that the “Database name” field should be the same as that used for the quality control step above. Once done, you can press the “Test connection” button to confirm that the settings are correct. The setting “Table Prefix” should be changed to a different value such that a new table is created and used for this step.
  vi. Click the ‘View output settings’ button at the top-left of the interface. In the settings panels, select an appropriate ‘Default output folder’ where the analysis data will be saved.
  vii. Optional: For smaller number of images (e.g., < 1000) or instances when a MySQL database is not available, the *ExportToSpreadsheet* module may be used to export the collected measurements to a comma-delimited file (CSV) format. Add this module by selecting *Edit > Add Module > File Processing > ExportToSpreadsheet* from the CellProfiler main menu, and position it as the last module in the pipeline by using the “^” or “v” buttons beneath the pipeline. Disable the *ExportToDatabase* module by clicking the green checkmark next to the module name; the green checkmark will then be grayed out to indicate its status. Select the *ExportToSpreadsheet* module, select “No” for “Export all Measurement types?”, and then select “Image” for the “Data to export” drop-down drop that appears. Click the “Add another data set” button and select “Nuclei” for the “Data to export” drop-down drop that appears. Click the “Add another data set” button again and select “Cells” for the “Data to export” drop-down drop that appears, and select “Yes” for the “Combine these object measurements with those of the previous object?” setting. Click the “Add another data set” button again and select “Cytoplasm” for the “Data to export” drop-down drop that appears, and select “Yes” for the “Combine these object measurements with those of the previous object?” setting. This will place two CSV files, containing the per-image and per-cell data, in the Default output folder.
  viii. Save the current settings to a project (.cpproj) file containing the pipeline, the list of input images and the output location by selecting *File > Save Project*. Enter the desired project filename in the dialog box that appears.
  ix. Use CellProfiler’s Test mode functionality (accessible from the “Test” menu item) to carry out analysis and visually inspect results from a small sample of images from across the experiment for accuracy of nuclei and cell body identification. Adjust image analysis pipeline parameters within CellProfiler as needed. The CellProfiler website contains resources and tutorials on how to optimize an image analysis pipeline. The Anticipated Results section outlines the expected nuclei and cell identification quality. **Critical step:** Because capturing subtle phenotypes is important for profiling, accurate nuclei and cell body identification is essential for success. Examine the outputs of *IdentifyPrimaryObjects* and *IdentifySecondaryObjects* for a few images to make sure the boundaries generally match expectations. Under the “Test” menu item, there are options for selecting sites for examination. We recommend either randomly sampling images for inspection (via “Random Image set”) and/or selecting specific sites (via “Choose Image Set”) from negative control wells or specific treatment locations from the plates. The rationale is to check a wide variety of treatment-induced phenotypes to ensure that the pipeline will generate accurate results. **(TROUBLESHOOTING)**
  x. Press the ‘Analyze Images’ button at the bottom-left of the interface. A progress bar in the bottom-left will indicate the estimated time of completion. **Critical step:** This step assumes that you will be running the image analysis pipeline locally on your computer, which generally is only recommended for experiments with <1000 fields of view. If your institution has a shared high-performance computing cluster, we recommend executing the pipeline on the cluster as a batch process, i.e., a series of smaller processes entered at the command line; this will result in much more efficient processing. Enlist the help of your institution’s IT department to find out whether this is an option and what resources are available. If so, carry out optional Step (d), describing modifications to the pipeline to run it as a batch process.
  xi. The pipeline will identify the nuclei from the Hoechst-stained image (referred to as “DNA” in CellProfiler), use the nuclei to guide identification of the cell boundaries using the SYTO 14-stained image (“RNA” in CellProfiler), and then use both of these features to identify the cytoplasm. The pipeline then measures the morphology, intensity, texture, and adjacency statistics of the nuclei, cell body and cytoplasm, and outputs the results to a MySQL database. See Supplemental Note 2 for the image features measured for each cell.
d. Configuring pipelines for batch processing on a computer cluster (optional)
  i. We recommend using a computing cluster for analyzing Cell Painting experiments to speed processing, especially for experiments with >1000 fields of view. The typical batch processing workflow is to distribute smaller subsets of the acquired images to run on individual computing nodes. Each subset is run using CellProfiler in “headless” mode, i.e., from the command line without the user interface. The headless runs are executed in parallel, with a concomitant decrease in overall processing time. **Critical step:** Carrying out this step requires significant setup effort and will probably require enlisting the help of your IT department. Please refer to the Supplementary Methods 1 or to our GitHub webpage https://github.com/CellProfiler/CellProfiler/wiki/Adapting-CellProfiler-to-a-LIMS-environment for more details.
  ii. Insert the *CreateBatchFiles* module into the pipeline by pressing the ‘+’ button, and selecting the module from the “File Processing” category. Move this module to the end of the pipeline by selecting with your mouse and using the ‘^’ or ‘v’ buttons at the bottom-left of the interface.
  iii. Configure the *CreateBatchFiles* module by setting the ‘Local root path’ and ‘Cluster root path’ settings. If your computer mounts the file system differently than the cluster computers, *CreateBatchFiles* can replace the necessary parts of the paths to the image and output files. For instance, a Windows machine might access files images by mounting the file system using a drive letter, e.g., C:\your_data\images and the cluster computers access the same file system using /server_name/your_name/your_data/images. In this case, the local root path is C:\ and the cluster root path is /server_name/your_name. You can press the ‘Check paths’ button to confirm that the path mapping is correct.
  iv. Press the ‘Analyze Images’ button at the bottom-left of the interface.
  v. The end result of this step will be a ‘Batch_data.h5’ (HDF5 format) file. This file contains the pipeline plus all information needed to run on the cluster.
  vi. This file will be used as input to CellProfiler on the command line, in order for CellProfiler to run in “headless” mode on the cluster. There are a number of command line arguments to CellProfiler that allow customization of the input and output folder locations, as well as which images are to be processed on a given computing node. Enlist an IT specialist to specify the mechanism for sending out the individual CellProfiler processes to the computing cluster nodes. Please refer to the Supplementary Methods 1 or to our GitHub webpage https://github.com/CellProfiler/CellProfiler/wiki/Adapting-CellProfiler-to-a-LIMS-environment for more details.

### 5. Normalize morphological features across plates

**Timing: < 5 min per 384-well plate**

a. The extracted features need to be normalized to compensate for variations across plates. For each feature, compute the median and median absolute deviation for all reference cells within a plate. The reference cells need not be a perfect negative control but instead simply provide a baseline from which other treatments can be measured. In RNAi and overexpression experiments, we have found untreated cells to be an effective baseline for normalization. For chemical experiments, we have found DMSO-treated cells to be an effective baseline.
b. Normalize the feature values for all the cells (both treated and untreated) in the plate by subtracting the median and dividing by the median absolute deviation (MAD) □ 1.4826 (multiplying the MAD by this factor provides a good estimate of the standard deviation for normal distributions).
c. Exclude features having MAD = 0 in any plate, because when this is the case, all samples have the exact same value for that feature and thus the feature does not carry any sample-specific information.
d. Sample profiling scripts (written in Python) to perform the normalization are are provided and described in Supplementary Methods 2.

### 6. Create per-well profiles

**Timing: 1 h per 384-well plate**

a. There are a number of possible approaches to create per-well profiles from the individual cell measurements from each image/site within the well. We (Broad Institute) have published a comparison of several such methods for creating morphological profiles^46^. Here, we describe the approach of population-averaging profiles, which has been shown to be effective. For each well, compute the median for each of the *n* features across all the cells in the well. This produces an *n*-dimensional data vector per well.
b. (Optional) Use principal components analysis (PCA) to reduce the dimensionality of the data. To do this, collect all the *n*-dimensional data vectors corresponding to all the *k* wells in the experiment, and produce an *n* x k dimensional data matrix. Perform PCA on this data matrix to obtain a lower dimensional representation of the per-well profiles. For instance, in one of our recent papers (Broad Institute)^30^, the dimensionality of the data vectors was reduced from 1301 to 205, while preserving 99% of the variance in the data.
c. Sample profiling scripts (written in Python) to create the per-well profiles are provided and described in Supplementary Methods 2.

### 7. Data analysis

**Timing: variable**

a. The per-well profiles created can be used to analyze patterns in the data. How to do so is an area of active research and is customized to the biological question at hand.
b. For example, morphological profiles were used to discover compounds that induce similar phenotypes using clustering^29^, to identify compound sets with high rates of activity and diverse biological performance in combination with high-throughput gene-expression profiles^34^, and to determine the dominance of seed-sequence driven off-target effects in RNAi-induced gene knockdown studies^30^. See the above cited publications for example analyses.

## TIMING

### Step 1, Cell culture

It typically takes about 2-3 days for the days for the cells to reach appropriate confluency and siRNAs to achieve appropriate knockdown depending on cell type and growth conditions. Harvesting the cells (steps 1e-1g) takes 30 minutes and seeding the cells (step 1h) takes 20 minutes. Optional transfection of siRNA takes one hour, including reagent preparation (steps 1d and 1i). After seeding the cells, we culture the cells for 2-4 days before staining (steps 1j-1n).

### Step 2, Staining and fixation

Approximately 2.5 - 3 hours including reagent preparation. The total timing will vary depending the number of plates in the experiment and the automation available. We have found that up to 12 plates can be simultaneously fixed and stained as one batch in this span of time, although we recommend no more than 4 - 5 plates per batch due to the increased likelihood of sample preparation error by the researcher.

### Step 3, Automated image acquisition

About 3.5 hours per 384-well plate, for 9 fields-of-view per well and the exposure times given in Table 1. The total time may vary depending on the number of sites imaged per plate and exposure time for each channel.

### Step 4, Morphological image feature extraction from microscopy data

Approximately 10 minutes per plate for CellProfiler to scan the inputs folder(s) after manually drag/dropping the needed images into the CellProfiler interface. The pipeline execution time will depend on the computing setup; run times on a single compute node of 20 sec (illumination correction), 30 sec (quality control), and 10 min (analysis) per field of view are typical. Performing the quality control workflow using CellProfiler Analyst takes approximately 4 hrs of hands-on time, though this time can be significantly shortened if cutoffs are re-used from experiment to experiment.

### Step 5, Normalize morphological features across plates

Under five minutes of processing time per 384-well plate. Additional time is required to set up data access infrastructure prior to this step.

### Step 6, Create per-well profiles

Up to an hour of processing time per 384-well plate, depending on data access methods.

### Step 7, Data analysis

Approximately one hour for basic analysis of replicate quality and signature strength. Time for additional analysis varies significantly depending on the problem at hand.

## TROUBLESHOOTING

| Step | Problem | Possible reason | Solution |
| --- | --- | --- | --- |
| 3c | The images contain bright, slender or punctate artifacts that appear in multiple wells, across multiple channels. Too many of these artifacts can adversely affect nuclei and cell body identification and measurement. | The washing reagents are contaminated with fibers, e.g. from clothing or dust. | Filter the washing solutions and diluents before use. Prepare plates in a clean, dust-minimal environment. |
| 4c(ix) | The identified nuclei or cell bodies do not reflect the actual boundaries of the stained nuclei or cells in the image. | The settings in the *IdentifyPrimaryObjects* or *IdentifySecondaryObjects* modules (for nuclei and cell identification, respectively) were optimized for U2OS cells imaged on a particular microscope at a particular magnification, and may be inappropriate for different experimental conditions. | Cell lines with different morphological features may require additional optimization of the pipeline identification modules. Visual inspection is needed to confirm that the settings conform to expected results. If you encounter difficulties in adjusting the pipeline settings for this task, we recommend consulting the moderated forum at http://www.cellprofiler.org/forum/ for assistance. |

## ANTICIPATED RESULTS

The automated imaging protocol will produce a large number of acquired images in 16-bit TIF format. The total number of images generated equals (number of samples tested) × (number of sites imaged per well) × (5 channels imaged). Results from a typical Cell Painting experiment (a small molecule study using A549 cells) are shown for a negative control DMSO well and digitoxin-treated well (Figure 2). An example image data set for an RNAi Cell Painting study of U2OS cells is supplied in Supplementary Note 1.

**Figure 2:**
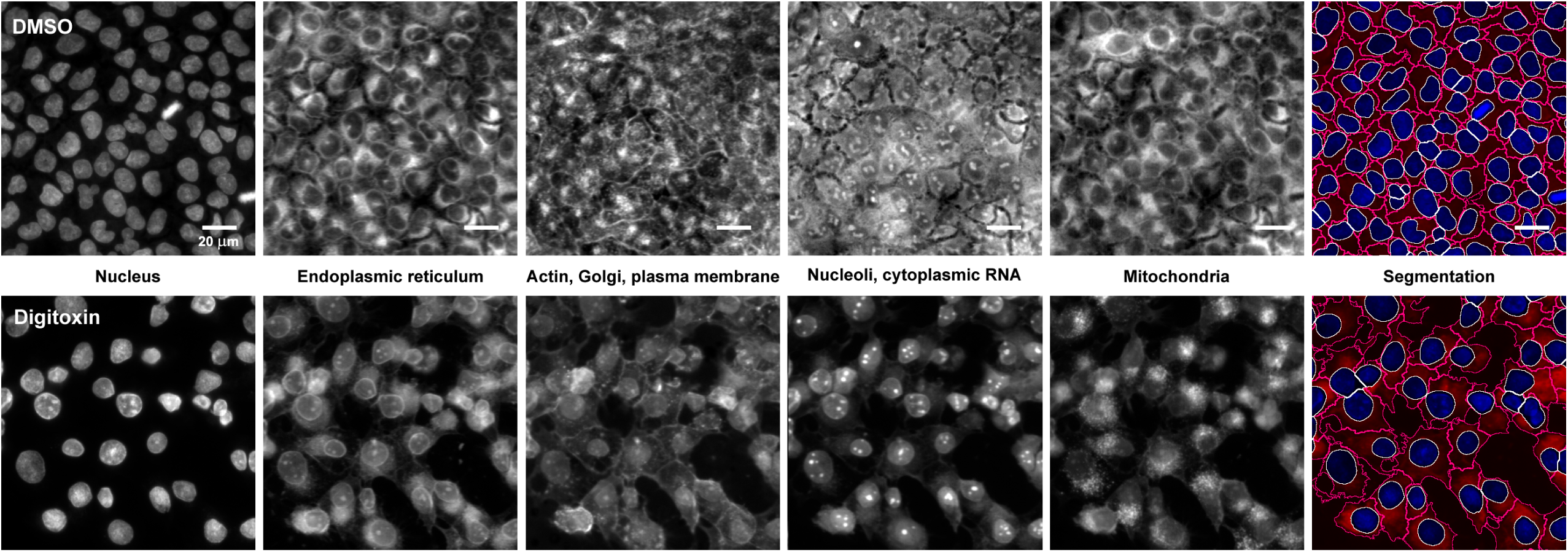
Sample images from a small molecule Cell Painting experiment using A549 cells. Images are shown from a DMSO (negative control) well (top row) and a digitoxin-treated well (bottom row). The first five columns display the five channels imaged in the Cell Painting assay protocol, while the last column illustrates a merged image of the DNA stain (blue) and the nucleolar/cytoplasmic RNA stain (red), with the identified (i.e, segmented) nuclei and cell body outlines resulting from image analysis overlaid in white and magenta, respectively. See Table 1 for details about the stains and channels imaged. Each panel is 6% of the full image; the total field of view is 9 images in a 3 × 3 grid, representing ~52% of the well area.

In addition, the illumination correction pipeline will yield five illumination correction images per plate, one for each channel; an example set of illumination correction images is provided as Supplemental Data 3. The quality control pipeline will produce a set of numerical measurements extracted at the image level, and export them to a MySQL database. These measurements can optionally be used for removing images that are unacceptable for further processing due to focal blur or saturation artifacts.

The image analysis pipeline will produce several outputs. Generally, the pipeline is not configured to save any processed images (to conserve data storage space) but the *SaveImages* module can be used for this purpose if desired, e.g., for saving outlines such as those in the last column of Figure 2. The pipeline also produces the raw numerical image features extracted from the cell images, which are deposited to a MySQL database. The database contains one row for each cell in each image, and ~1000 columns containing the values for the different morphological features that have been measured for that cell. A sample set of morphological profiling data is provided as Supplementary Data 4.

The quality of the extracted image features and downstream profiling will depend on accurate nuclei and cell body segmentation. The last column of Figure 2 contains overlays of the nuclei and cell body identification (i.e, segmentation) to highlight the differences in cellular morphology between the two treatments. First, the nuclei are identified from the Hoechst image because it is a high-contrast stain for a well-separated organelle; subsequently, the nucleus along with an appropriate channel is used to delineate the cell body^47^. We have found the SYTO 14 image is the most amenable for finding cell edges, as it has fairly distinct boundaries between touching cells. We have found that very little adjustment of the analysis pipeline is needed to achieve good quality data, even across multiple cell lines. Even so, it is important to understand the key segmentation parameters of the pipeline in order to optimize the output if needed (see Troubleshooting note for Step 3, “Automated image acquisition”).

After running the profiling scripts (Supplementary Methods 2, written in Python) to normalize the image features across plates and create the per-well profiles, the output will be a morphological profile file in comma-delimited format (Supplementary Data 5, CSV). Each row of this file represents data vector for an individual plate and well, with each column containing the median for each of the ~1000 image features across all the cells in that well.

## SUPPLEMENTARY INFORMATION

- Supplementary Data 1: A ZIP file containing a sample ImageXpress plate acquisition settings file https://github.com/carpenterlab/2016_bray_natprot/raw/master/supplementary_files/ImageXpress_CellPainting_plate_acqusition_settings.zip.
- Supplementary Data 2: A ZIP file containing the CellProfiler pipelines described in the protocol: Illumination correction, quality control, and feature extraction. Pipelines were created using CellProfiler 2.1.1. https://github.com/carpenterlab/2016_bray_natprot/raw/master/supplementary_files/cell_painting_pipelines.zip
- Supplementary Data 3: A ZIP file containing example illumination correction images generated by the illumination correction pipeline for the images provided by Supplementary Note 1. https://github.com/carpenterlab/2016_bray_natprot/raw/master/supplementary_files/illumination_correction_images.zip
- Supplementary Data 4: Files containing an example morphological profiling dataset generated with image data supplied in Supplementary Note 1. https://github.com/carpenterlab/2016_bray_natprot
- Supplementary Data 5: A ZIP file containing a CSV of per-well profiles generated by running the steps in Supplementary Methods 2 on the dataset in Supplementary Data 4. http://pubs.broadinstitute.org/bray_natprot_2016/suppl/online/profiles.zip
- Supplementary Note 1: Image data from an RNAi Cell Painting knockdown experiment applied to U2OS cells may be found at https://www.broadinstitute.org/bbbc/BBBC025/
- Supplementary Note 2: A PDF listing of per-cell image features generated by CellProfiler. https://github.com/carpenterlab/2016_bray_natprot/raw/master/supplementary_files/cellprofiler_feature_listing.pdf
- Supplementary Methods 1: A screen capture PDF of GitHub wiki webpage: https://github.com/CellProfiler/CellProfiler/wiki/Adapting-CellProfiler-to-a-LIMS-environment.
- Supplementary Methods 2: Steps to produce per-well morphological profiles from single cell measurements. https://github.com/carpenterlab/2016_bray_natprot/raw/master/supplementary_files/profiling_methods.pdf

## AUTHOR CONTRIBUTION STATEMENTS

All authors contributed to writing this protocol. CTD, HH, and SMG contributed to describing the benchwork aspects of the protocol. CH, MKA, MAB and CTD contributed to updates to the experimental design. MAB and SS contributed most heavily to describing the computational aspects of the protocol. AEC, CCG, BB, and SS contributed most heavily to describing the rationale and experimental design.

## ACKNOWLEDGMENTS

Research reported in this publication was supported in part by NIH R44 TR001197 (CCG), NSF RIG DBI 1119830 (MAB), NIH R01 GM089652 (AEC) and NSF CAREER DBI 1148823 (AEC). The RNAi Cell Painting knockdown experiment used in this publication was previously published^30^ and was supported in part by the Slim Initiative for Genomic Medicine, a project funded by the Carlos Slim Foundation in Mexico. The authors thank the original developers of earlier versions of the protocol, who are the authors of the original paper describing the assay^29^; these authors are: Sigrun M. Gustafsdottir, Vebjorn Ljosa, Katherine L. Sokolnicki, J. Anthony Wilson, Deepika Walpita, Melissa M. Kemp, Kathleen Petri Seiler, Hyman A. Carrel, Todd R. Golub, Stuart L. Schreiber, Paul A. Clemons, Anne E. Carpenter, Alykhan F. Shamji. For enabling this work, Recursion Pharmaceuticals thanks the University of Utah Core Facilities (John Phillips), specifically the Drug Discovery Core (Bai Luo) and the Fluorescent Imaging Core (Chris Rodesch). The Broad thanks members of the Broad Institute’s Center for the Development of Therapeutics, especially Thomas P Hasaka for technical assistance. We also thank Alison Kozol for proofreading the equipment and reagents and for testing the image analysis workflow, and Alice Berger and Xiaoyun Wu for offering helpful comments and suggestions during manuscript preparation.

